# Generating Templates and Growth Charts for School-Aged Brain Development

**DOI:** 10.1101/747352

**Authors:** Hao-Ming Dong, F. Xavier Castellanos, Ning Yang, Zhe Zhang, Ye He, Lei Zhang, Ting Xu, Avram J. Holmes, B.T. Thomas Yeo, Feiyan Chen, Bin Wang, Christian Beckmann, Tonya White, Olaf Sporns, Jiang Qiu, Tingyong Feng, Antao Chen, Xun Liu, Xu Chen, Xuchu Weng, Michael P. Milham, Xi-Nian Zuo

## Abstract

Standard brain templates and growth charts provide an invaluable resource for basic science research, with the eventual goal of contributing to the clinical care of neuropsychiatric conditions. Here, we report on a protocol to generate MRI brain templates in children and adolescents at one-year intervals from 6-to-18 years of age, with their corresponding growth charts, using a large-scale neuroimaging data resource (948 brain images from China and United States). To assure that the brain templates and growth charts are reliable and accurate, we developed a refined pipeline consisting of template construction, image registration, brain area labeling and growth chart modeling. The pipeline comprises multiple modular workflows that can be used for multiple applications. In our approach, population- and age-specific templates were first constructed to avoid systemic bias in registration. Brain areas were then labeled based on the appropriate templates, and their morphological metrics were extracted for modeling associated growth curves. We implemented warp cost as a function of age differences between individual brains and template brains. A strong U-shaped cost function was revealed, indicating larger age differences are associated with greater registration errors. This validates the necessity of age-specific reference templates in pediatric brain imaging studies. Growth chart analyses revealed preferential shape differences between Chinese and US samples in lateral frontal and parietal areas, aspects of cortex which are most variable across individuals with regard to structure and function as well as associated behavioral performance. This growth distinction is largely driven by neurodevelopmental differences between Chinese and US age-specific brain templates. The pipeline together with the brain templates and charts are publicly available and integrated into the Connectome Computation System.

## Introduction

Growth charts are an invaluable resource for enhancing public health. They are essential for screening the developmental status of individuals and monitoring their abnormal growth as an early detection tool^1^. Deviations from normative age-expected values are used to trigger evaluations for underlying abnormal factors, which can provide important developmental information to clinicians and parents. Extending this approach to the evaluation of an individual’s neurodevelopmental status has been impeded by the lack of reliable growth charts for the human brain. Magnetic resonance imaging (MRI) is increasingly being employed to map human brain development. Anatomical MRI (aMRI) can capture developmental changes of brain morphology^2, 3^, which comprise full-brain geometrical transformations (e.g., cortical thinning and surface expansion)^4, 5^. For example, changes in cortical thinning trajectories have been linked with inter-individual differences in IQ in children and adolescents^6^. Such developmental effects in brain structure have also been shown to be detectable across adulthood^7^ and are supported by brain network studies using diffusion-weighted (dMRI) and resting-state functional (rfMRI) imaging methods^8, 9^, providing the framework for quantifying multimodal brain development at the population level^10, 11^. Although sparse, efforts to translate developmental trajectories into growth charts have begun to be initiated for neuropsychiatric conditions^12–14^, which are believed to have abnormal neurodevelopmental origins^15, 16^.

Despite the promise of developmental population neuroscience, a number of key issues must be addressed prior to establishing brain growth charts for clinical use. First, reliability of MRI-based measurements must meet clinical standards on measurements of individual differences^17–19^. Core anatomic MRI measures (e.g., volume, cortical thickness, surface area) currently meet this standard^20^, but most dMRI and rfMRI measures do not due to multiple confounds and substantial random error^21, 22^. This suggests aMRI-derived measures could provide the bases for developing reliable imaging markers of clinically useful growth charts. Their high reliability makes it possible to attain highly valid charts, though of course, does not guarantee this^23^. Second, MRI samples of brain development cohorts for building growth charts are currently limited. Large-scale brain development cohorts are fundamental for charting growth^24–26^, but unfortunately obtaining longitudinal assessments across multiple centers with the same protocols is rare^27, 28^. Previous studies have demonstrated the necessity of creating growth charts for height, weight and head circumference for specific populations or countries^29^, which is likely even more important for brain growth charts, given the neurodevelopmental diversity likely arising from differences in ethnicity and culture. Finally, detection of differences could be biased by using inappropriate analyses performed at the population level. For example, neuroimaging studies have already documented biases from using generic brain templates for multiple age ranges^30–35^, indicating the need of a full range of age-specific brain templates across school age (6-18 years). Despite the public health importance of creating normative charts^12, 36–39^, a protocol addressing these issues to generate brain templates and growth charts for school-age development is lacking.

This protocol was designed to begin to fill this gap. It consists of an integrative pipeline for generating brain templates and growth charts of children and adolescents. Volumetric measurements were quantified with aMRI of 674 school-age brains from two accelerated longitudinal cohorts with the same experimental design obtained in the United States (Enhanced Nathan Kline Institute Rockland Sample - eNKI sample)^40^ and China (Chinese Color Nest Project - CCNP)^10, 41^, respectively. Standard brain templates were constructed annually for each year of age and serve as a field-wide resource for generating growth charts on morphological development of brain tissues, lobes and networks. These brain templates and growth charts were validated across two cultures to offer an initial normative reference for studies of school-age brain development.

## Development of the protocol

Construction of reliable brain grow charts at the area-level relies heavily on the accurate localization of brain areas across individuals, i.e., MRI image registration. Registration is commonly used to automatically label individual images from atlases defined on standard brain templates. Previous studies^33, 34, 42^ have shown two factors that account for the most variance during template registration, ethnicity and age. Ethnicity plays a critical role in shaping brain morphology^43^. For instance, significant volumetric differences were observed between Chinese and Caucasian adult brain templates^33, 42^, indicating a rounder global shape and shorter axial distance in Chinese adults. Dynamic neurodevelopmental factors affect brain maturation, suggesting such brain morphological differences should also be observable during childhood or adolescence. Therefore, creating a custom brain template from a homogenous population has been strongly recommended to improve registration performance^44^. However, the desirability of population- and age-specific brain templates for modeling growth charts has not been prioritized.

In the single exception, group differences in registration errors relating to ethnic and developmental factors were tested^34^. However, the utility and generalizability of their templates was limited by the small sample size (n=138) from a single imaging site using relatively broad age intervals (2 years). Moreover, the relationship between registration errors and age in pediatric samples has yet to be examined and quantified.

Thus, to improve the accuracy of brain growth charts, we established a protocol, i.e., a pipeline consisting of brain template construction, image registration, regional area labeling and growth chart modeling. In the pipeline, two population- and age-specific templates (Figure 1) were first constructed to avoid systemic bias in registration (Institute of Psychology, Chinese Academy of sciences (IPCAS) and Nathan Kline Institute (NKI) brain templates), then brain areas were automatically labeled based on the age- and ethnicity-matched templates, and finally their morphological metrics were extracted for modeling growth charts.

**Figure 1.**
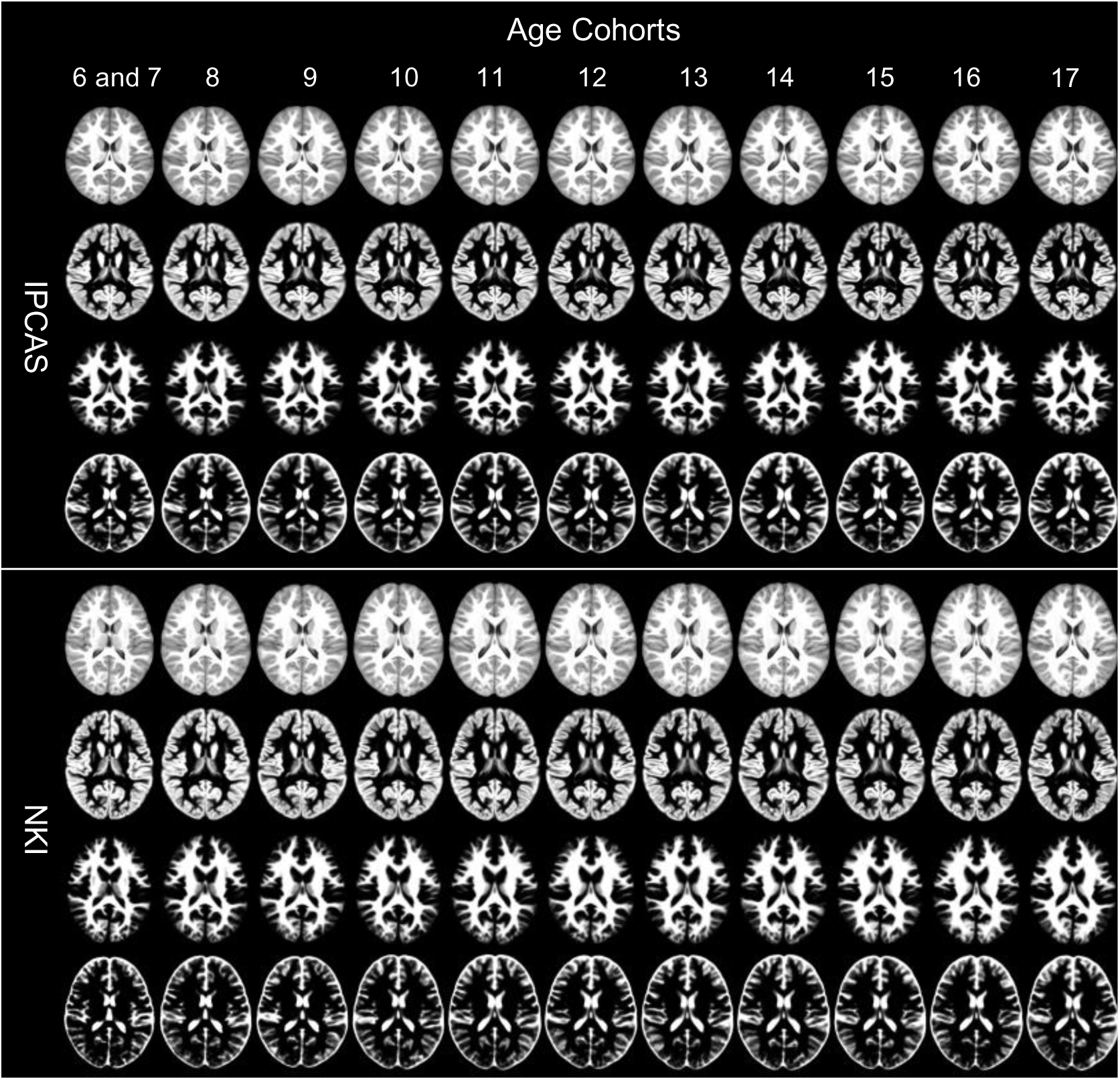
Axial slides of brain and tissue probability templates. IPCAS (up 4 rows) and NKI (bottom 4 rows) templates with one-year interval (except age 6 and age 7) are listed from top to bottom in sequence of templates of brain, gray matter probability, white matter probability and cerebrospinal fluid probability.

## Comparison with other methods

Growth charts built with existing big datasets provide clear evidence that it is necessary to estimate brain morphological properties as well as their corresponding brain templates within specific age groups. This is especially true for pediatric neuroimaging studies of the school-age population, in which there are considerable changes in brain morphology. Both cross-sectional and longitudinal applications of growth charts are facilitated by choosing proper templates, where the typical value is delivering valid atlas information, i.e., defining regional areas in individual brains.

Previous studies have demonstrated that different ages and ethnicities increase the deformation costs associated with morphing anatomical regions between individual brains, which if done poorly, can result in mismatches in brain segmentation tissue profiles^33, 34^. This is supported by our findings that even with an identical brain atlas, morphological metrics can differ substantially when registered to different brain templates. Ideally, these metrics should be identical. In practice, the method with lower registration errors or costs is preferred.

As demonstrated in Figure 2, we observed that registration costs were related to national origin and stage of development. This was particularly well illustrated by using age-specific brain templates for the longitudinal CCNP and eNKI samples to model growth charts of brain volume (Figure 3), which were similar (peaking at 12-13 years of age but differing in specific details) to the inverted-U shaped curves observed in previous studies^45, 46^. The eNKI sample exhibited larger brain volumes and more accelerated increases during childhood than the CCNP sample. Differences in such a fundamental morphological characteristic may lead to increases of registration errors related to age and ethnic differences (Figure 2). Using an independent validation sample (n=84, 7-12 years, Chinese), we compared the deformation costs of registering the individual brains to the IPCAS and NKI templates across different ages. As expected, the NKI brain templates resulted in greater image deformations than the IPCAS templates (red versus blue). Beyond this observation, registration deformations associated with age-matched brain templates were less than those of age-mismatched templates. As revealed in Figure 2, the smallest deformations occurred when sources and targets of brain registration were approximately age matched, with a little deviation to negative matching ages referring to the templates built with younger (both CCNP and eNKI) samples.

**Figure 2.**
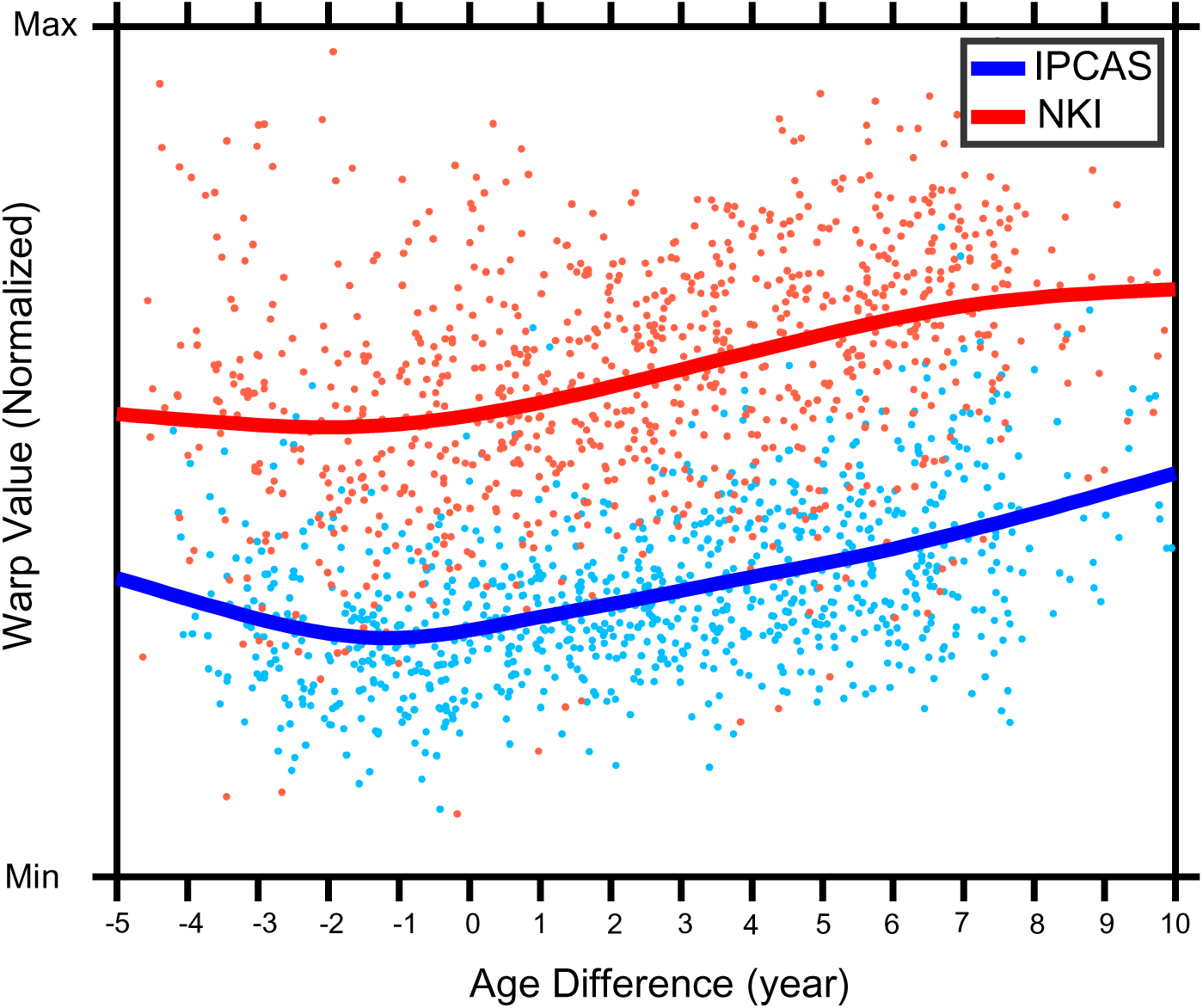
Deformation cost as a function of age difference. GAMLSS fits on the deformation cost (normalized warp values) with age differences between individual validation brains and template brains. Blue indicates the use of IPCAS pediatric templates while red indicates the use of eNKI pediatric templates for the registration.

**Figure 3.**
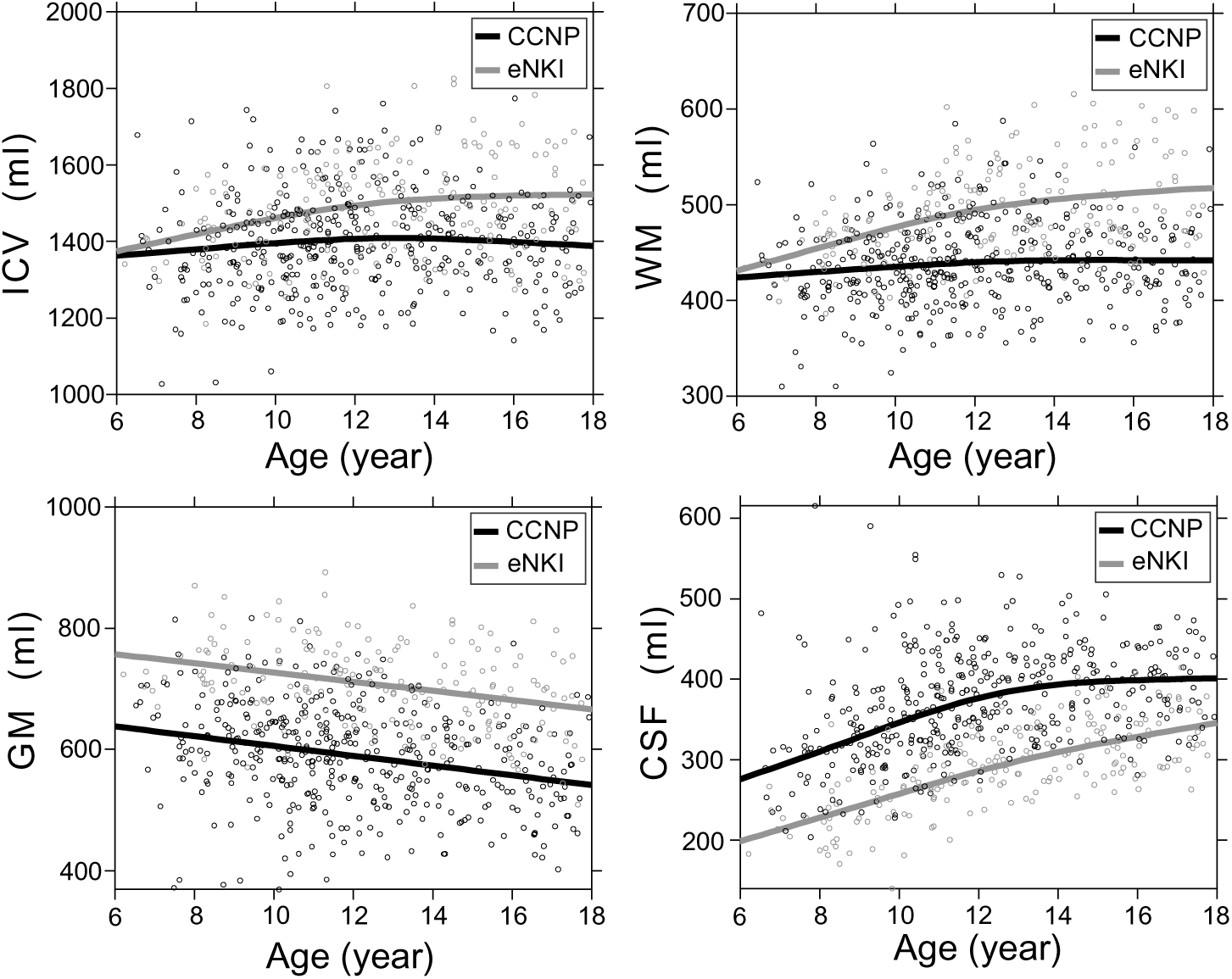
Growth charts of brain tissue volume. Growth charts of Intracranial Volume (ICV), Gray Matter (GM), White Matter (WM) and Cerebral Spinal Fluid (CSF). The black line is derived from CCNP samples while the gray line is based on eNKI samples.

The IPCAS pediatric templates were generated with much larger sample sizes and narrower intervals for longitudinal sampling than prior efforts. First, to the best of our knowledge, CCNP is the largest longitudinal MRI database of Chinese healthy school-age children. The number of scans (n = 674) is nearly 5 times and 13 times larger than previous studies by Xie et al.^34^ (n = 138) and Luo et al.^32^(n = 53), respectively. Second, the pediatric MRI images were collected from typically developing children recruited from primary and middle/high schools at three imaging sites, which are widely distributed in China, making the sample more representative of the Chinese healthy pediatric population than samples recruited from clinical sites.

For depicting brain developmental trajectories, results appear to be dramatically driven by the age-specific templates. For instance, we consider an area located in the right superior parietal gyrus (labeled as Parcel No.136 of the frontal-parietal network in Schaefer et al.^47^) (Figure 4). Its growth curve exhibited relatively distinct patterns between CCNP and eNKI samples when individual brains were registered to ethnic- and age-appropriate templates. However, growth patterns were inverted when individual brains were deformed to the mismatched ethnic-template. Specifically, the pattern of the growth charts was largely driven by the developmental changes of the employed brain templates. The observation that extraction of the areal metric largely depended on the target templates used for registration held generally across the whole brain.

**Figure 4.**
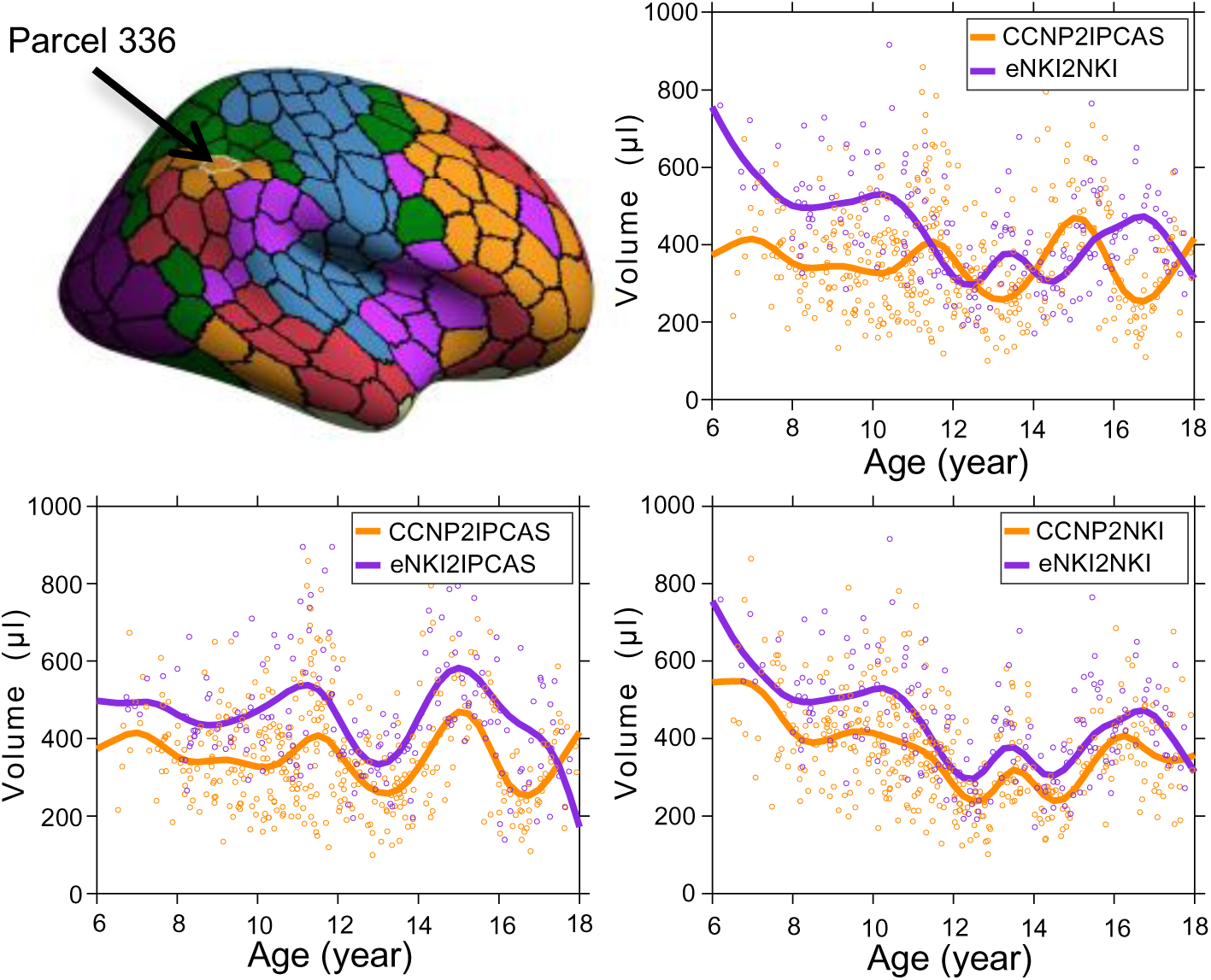
Template effects on growth charts of regional volume. The location of parcel 336 is shown in the upper left panel. Upper right panel shows the charts estimated with registrations using ethnic- and age-appropriate templates, panels at second row show charts estimated with registrations using ethnic-inappropriate templates. eNKI2IPCAS indicates eNKI samples registered to the IPCAS template while CCNP2NKI indicates CCNP samples registered to the NKI template. CCNP and eNKI are samples registered to ethnic- and age-appropriate templates.

Use of ethnic- and age-specific brain templates have not been used in previous developmental studies. This is partially because small samples are insufficient to construct such templates and few developmental studies have focused on such areal scales (small parcels)^47^. In the present protocol, we showed that for growth chart modeling, use of improper brain templates would unexpectedly and substantially distort the estimations of underlying morphological development, making conclusions questionable.

## Applications of the method

For a growing number of longitudinal neuroimaging cohort studies across the globe in recent years^24–28, 40, 41^, identification of typical developmental patterns is one of the primary research targets. The absence of a standard protocol impedes generalization between ethnic backgrounds, imaging settings and data processing procedures. This protocol was initially designed for generating validated and accurate growth charts but is not limited to only this application. Any MRI study that focuses on group-level information of individual brains would result in better precision by applying proper brain templates, especially for interracial and developmental research in which large morphological differences are expected across groups.

## Limitation and future work

Several limitations must be considered regarding the application of pediatric templates and the interpretation of growth charts. The proportion of males and females was balanced in most age groups except the 15-, 16- and 17-year-old age groups. Given previously reported sex differences in brain development^46, 48, 49^, constructing sex-specific templates in the future would be desirable. Many factors can affect the construction of pediatric growth charts, including the data preprocessing pipeline^48^, modeling methods^10^, and site effects^50^. Dynamic developmental trajectories might be confounded by image registration errors if inappropriate brain templates are employed. The construction of age-specific brain templates and developmental trajectories or growth charts should be performed in tandem. Finally, the age intervals used to define templates in the present studies were defined provisionally due to the lack of more detailed evidence on brain development. Nonetheless, the age-specific brain templates generated in the current study can facilitate the estimation of more precise changes in human brain morphology during development. Regional volume was employed in this protocol to demonstrate age and ethnicity effects on brain templates and growth charts. It is an interesting topic to investigate how such effects can be generalized to other metrics of human brain morphometry (see reference^51^ for a review).

## Overview of the procedure

We developed a pipeline to construct age-specific brain templates and brain growth charts together. Specifically, using a large neuroimaging dataset of Chinese pediatric brain images, we demonstrate for the first time that greater age mismatching of templates introduces larger registration deformations. Further, these age-specific templates can improve the accuracy of image registration between individual pediatric structural brain images, thereby facilitating more reliable and accurate human brain mapping studies in healthy and clinical pediatric populations. By modeling growth charts, we found that differences across western and eastern samples were decreased when examined at large-scale levels, including tissue classes of brain lobe volumes. At more fine-grained levels of spatial resolution, ethnic differences in cortical surface area indices became markers, particularly in association cortex, which exhibits greater flexibility, morphological variability and hemispheric asymmetry^52^.

## Experimental Design

To chart brain growth models, we developed a standard pipeline consisting of customized brain template construction, robust imaging registration and growth chart estimation. Ethnicity and age are the two major variables addressed in this work. For the first two of these steps, we examined a 2 (ethnic levels, CCNP vs. eNKI) *×* 11 (age levels) within-subject design to test template effects in registration. The 11 age levels ranged from 6 to 17 years old; with ages 6 and 7 combined into one group due to increase sample size (the sample size per age group can be found in **Materials**). This generated 22 ethnicity- and age-specific templates. Individual brain images from a validation dataset were then registered to these 22 templates, with 22 corresponding registration deformations calculated for each subject. Previous studies applied group-level comparisons in which registrations were divided into appropriate and inappropriate groups for estimating the template effects, with paired T tests or variance analysis performed to assess ethnicity differences in template registration^31^. We believe that between-group comparisons are insufficient for revealing age effects in registration cost, especially for age-ranges spanning from childhood to adolescence. Hence, we used continuous age differences instead of a categorical age group variable to model developmental changes. Finally, two curves, corresponding to the ethnicity factors (CCNP vs. eNKI), with age difference as an independent variable and deformation value as the dependent variable, were modeled to show ethnicity and age effects.

For modeling the growth charts of brain morphological metrics, the ethnicity factor was considered as potentially a confounding variable. Due to the lack of ethnic-specific templates in the past, the MNI template has been usually utilized as the default. To test how an ethnicity-unspecific template affects morphological estimation, we performed a 2 *×* 2 mixed design with ethnicity as the between-group factor (CCNP vs. eNKI) and appropriateness as the within-group factor (brain images registered to ethnic-matched and ethnic-mismatched templates). Two registrations were performed for each participant. For a child from the CCNP sample, ethnicity-appropriate registration refers to using an IPCAS age-appropriate template while ethnicity-inappropriate refers to using an NKI age-appropriate template, and similarly for participants in the eNKI sample. The 400-unit areal parcellation (in MNI space)^47^ was extracted based on the above two registrations for each subject and their growth charts were modeled, generating four growth charts (gc) for each brain area: 1) CCNP-gc (CCNP samples registered to the IPCAS templates), 2) eNKI-gc (eNKI samples registered to the NKI template), 3) CCNP2NKI-gc (CCNP samples registered to the NKI template), 4) eNKI2IPCAS-gc (eNKI samples registered to the IPCAS template). We hypothesized that the former two charts would be more appropriate than the latter two charts. We calculated the similarities of the volume growth charts for each parcel and grouped local areas into seven large-scale networks^47^ (Figure 5).

**Figure 5.**
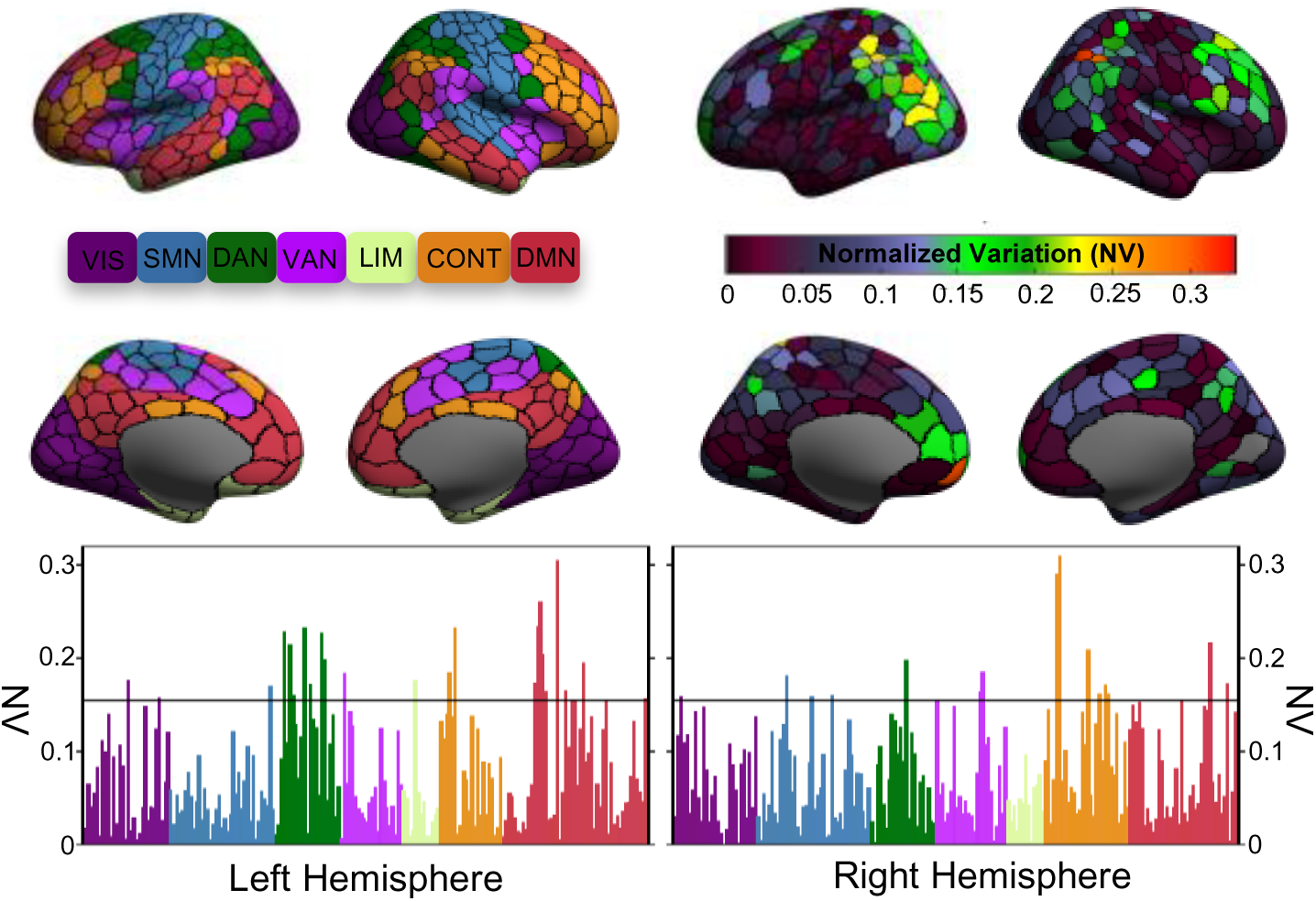
Similarities of brain growth charts between CCNP and eNKI samples. Upper left panel shows the 400 brain parcellation units, with parcels colored according to the Yeo2011 seven-network organization. Upper right panel shows similarities of the brain growth charts between CCNP and eNKI samples as measured by normalized variances (NV). The second row depicts the corresponding NV bar charts.

## Materials

### Equipment

#### Software

A computer with Linux or Unix environment or MAC OS pre-installed and with Bash shell scripting is required to run the algorithm.

Neuroimaging processing software included:

FSL (http://fsl.fmrib.ox.ac.uk/fsl/fslwiki/FslInstallation)53
ANTs (http://stnava.github.io/ANTs)54
volBrain (http://volbrain.upv.es)55

Statistical software R with the GAMLSS^56^ package installed for growth chart modeling.

### Template construction

#### Participants

MRI scans (n=774) were collected from 496 school-age (age range 6-18 years) typically developing children (TDC) of the Chinese Han population at three brain imaging sites. The final dataset passing quality control procedures consisted of the SWU413 sample^41^ (196 TDC scanned at the Faculty of Psychology, Southwest University in Chongqing), the SMU130 sample^57^ (130 TDC scanned at the First Hospital of Shanxi Medical University in Taiyuan) and the PKU131 sample^58–60^ (131 TDC obtained from the Beijing site of ADHD200 sample shared via the International Data-sharing Initiative). Specifically, the SWU413 dataset was acquired as a trial stage implementation of the developmental phase of the Chinese Color Nest Project (CCNP-SWU413)^10^, which is a five-year accelerated longitudinal study designed to delineate normative trajectories of brain development of Chinese children^41^. The age and sex distributions of overall MRI scans for the three samples are listed in Table 1.

**Table 1.** Age and sex composition in the three Chinese samples

| Age range | Chonqing site |  | Shanxi site |  | Beijing site |  | male | total |
| --- | --- | --- | --- | --- | --- | --- | --- | --- |
|  | male | total | male | total | male | total |  |  |
| **6.0-6.9** | 4 | 6 | 1 | 1 | 0 | 0 | 5 | 7 |
| **7.0-7.9** | 9 | 19 | 2 | 3 | 0 | 0 | 11 | 22 |
| **8.0-8.9** | 16 | 33 | 1 | 6 | 5 | 13 | 22 | 52 |
| **9.0-9.9** | 25 | 44 | 0 | 4 | 10 | 25 | 35 | 73 |
| **10.0-10.9** | 30 | 66 | 5 | 11 | 10 | 19 | 45 | 96 |
| **11.0-11.9** | 37 | 60 | 11 | 22 | 15 | 26 | 63 | 108 |
| **12.0-12.9** | 23 | 42 | 7 | 24 | 11 | 15 | 41 | 81 |
| **13.0-13.9** | 17 | 38 | 8 | 21 | 16 | 22 | 41 | 81 |
| **14.0-14.9** | 19 | 31 | 8 | 18 | 8 | 10 | 35 | 59 |
| **15.0-15.9** | 5 | 22 | 1 | 12 | 0 | 1 | 6 | 35 |
| **16.0-16.9** | 7 | 30 | 2 | 6 | 0 | 0 | 9 | 36 |
| **17.0-17.9** | 6 | 22 | 2 | 2 | 0 | 0 | 8 | 24 |
| **total** | 198 | 413 | 48 | 130 | 75 | 131 | 321 | 674 |
| **children** | 121 | 228 | 20 | 47 | 40 | 83 | 181 | 358 |
| **adolescents** | 77 | 185 | 28 | 83 | 35 | 48 | 140 | 316 |

For the enhanced NKI (eNKI) Rockland Sample^40^, a total of 561 scans were collected from 323 school-aged children. After the same quality control procedure applied for CCNP samples, a total of 190 scans from 133 TDC were included for our final analyses. Of note, CCNP and eNKI datasets both are accelerated longitudinal designs, were initially designed with matched age span and imaging resolution. Participants in the CCNP and eNKI sample who had a history of neurological or mental disorder, family history of such disorders, organic brain diseases, physical contraindication to MRI scanning, a total Child Behavior Checklist (CBCL) T-score higher than 70, or a Wechsler Intelligence Scale for Children IQ standard score lower than 80 were excluded.

CCNP and eNKI projects obtained the Institutional Review Board approval from IPCAS and NKI respectively. Written informed assent and consent were obtained from both participants and their parents/guardians. The details of the other samples can be found in previous reports^57–59^. According to the matched age and imaging resolution as well as the identical experimental design (Table 2 and Figure 6), both CCNP-SWU413 and eNKI samples were employed for the growth chart modeling. As few children were 6 (n=7) or 7 (n=22) years old, these two age groups were combined into a single group.

**Figure 6.**
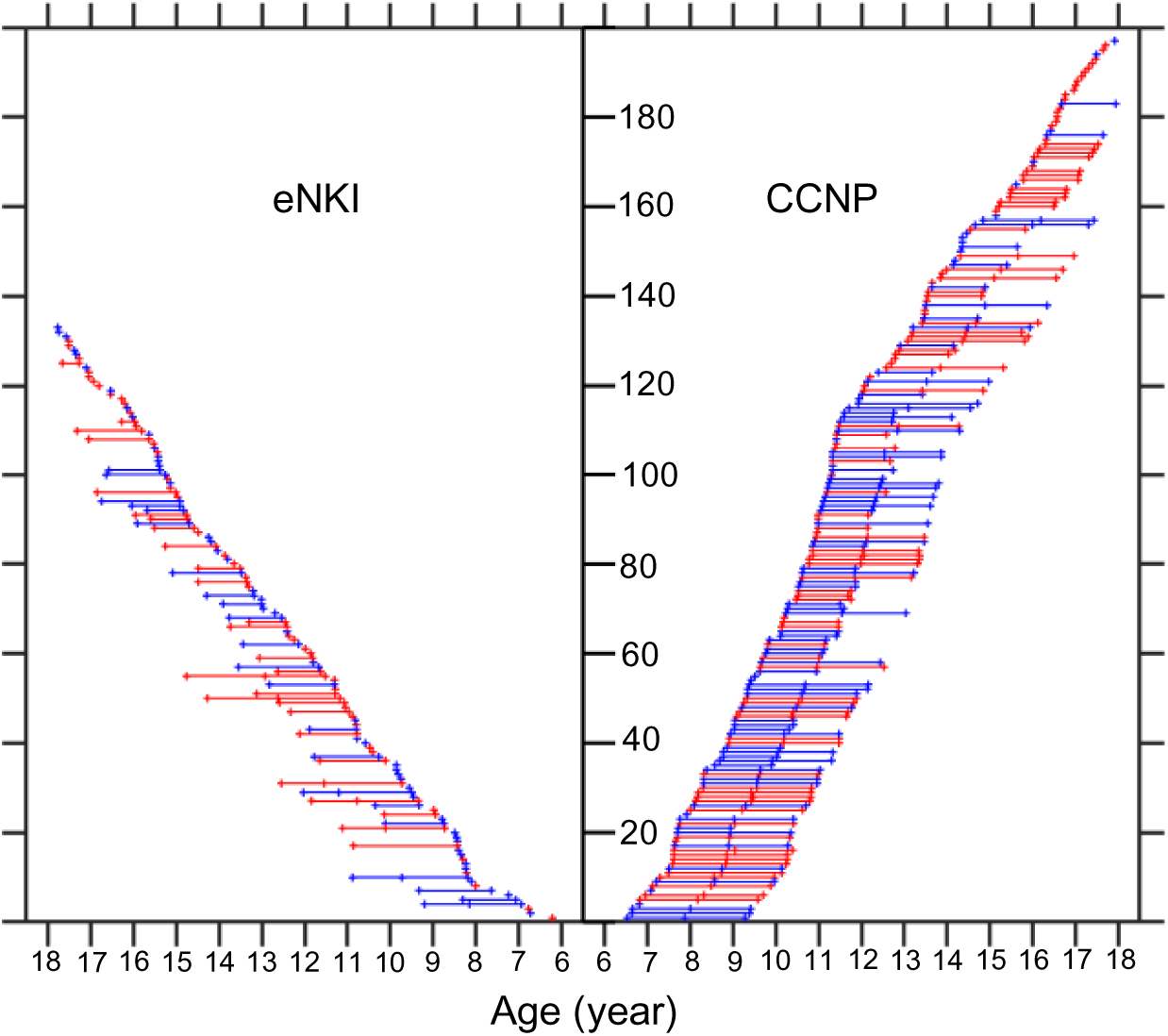
Age and sex distributions of CCNP and eNKI samples. Red indicates females while blue indicates males.

**Table 2.** Age and sex composition in CCNP and eNKI samples

| Age range | CCNP Samples |  | eNKI Sample |  |
| --- | --- | --- | --- | --- |
|  | male | total | male | total |
| **6.0-6.9** | 4 | 6 | 2 | 4 |
| **7.0-7.9** | 9 | 19 | 0 | 3 |
| **8.0-8.9** | 16 | 33 | 7 | 20 |
| **9.0-9.9** | 25 | 44 | 2 | 13 |
| **10.0-10.9** | 30 | 66 | 11 | 19 |
| **11.0-11.9** | 37 | 60 | 15 | 21 |
| **12.0-12.9** | 23 | 42 | 11 | 18 |
| **13.0-13.9** | 17 | 38 | 10 | 20 |
| **14.0-14.9** | 19 | 31 | 10 | 18 |
| **15.0-15.9** | 5 | 22 | 13 | 24 |
| **16.0-16.9** | 7 | 30 | 8 | 15 |
| **17.0-17.9** | 6 | 22 | 9 | 15 |
| **total** | 198 | 413 | 98 | 190 |
| **children** | 121 | 228 | 37 | 80 |
| **adolescents** | 77 | 185 | 61 | 110 |

### MRI scanning protocol

All data were acquired with Siemens Trio 3.0T scanners at all four imaging sites (see Table 3 for details of the scanning protocols at the Beijing site, Table 4 for details at Chongqing, Taiyuan, and Rockland sites). The scanning procedures across these sites can be found in previous publications^57–59^ and the FCP website (http://fcon_1000.projects.nitrc.org/indi/adhd200).

**Table 3.**
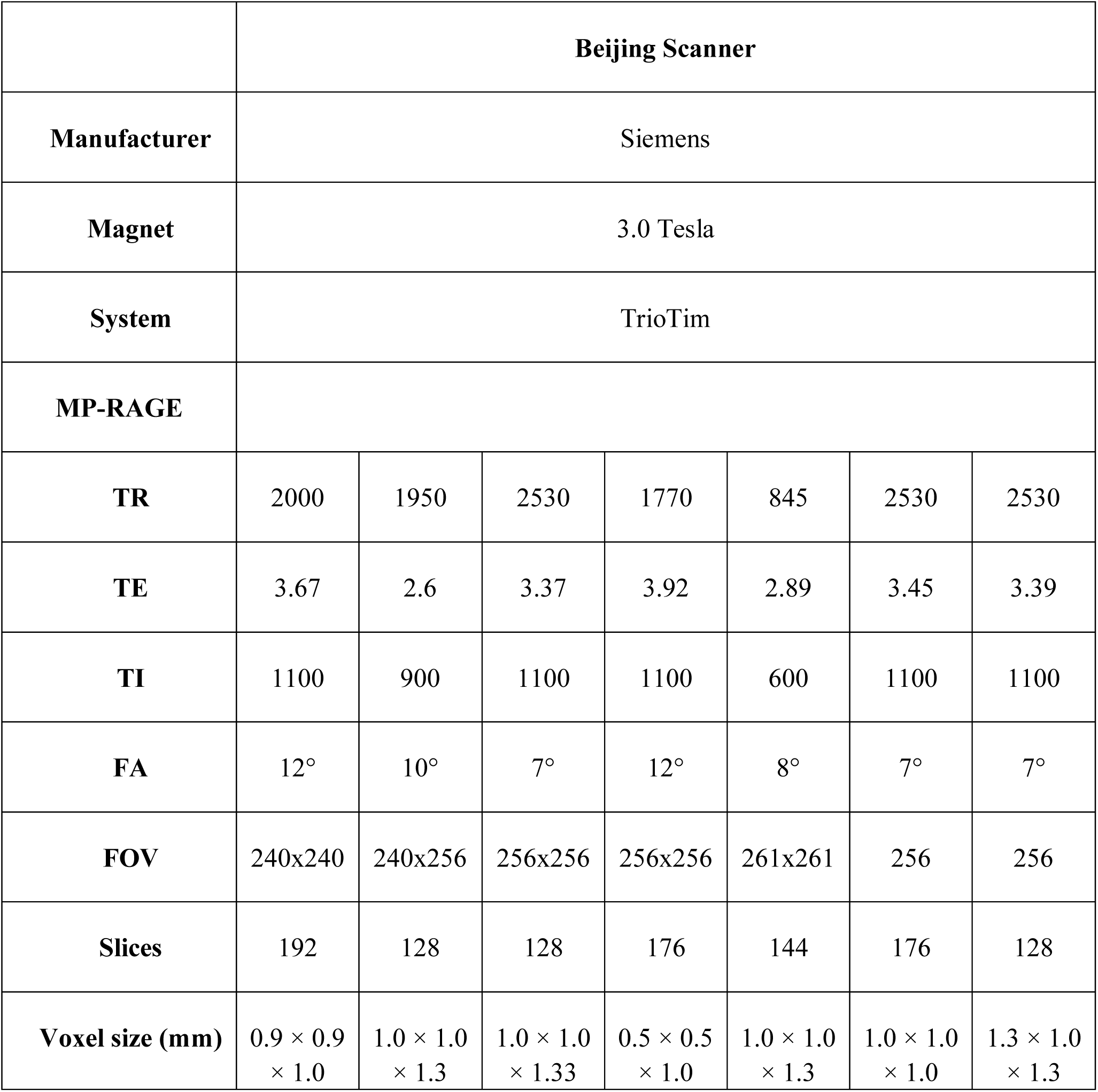
MRI scanning parameters at Beijing sites

**Table 4.**
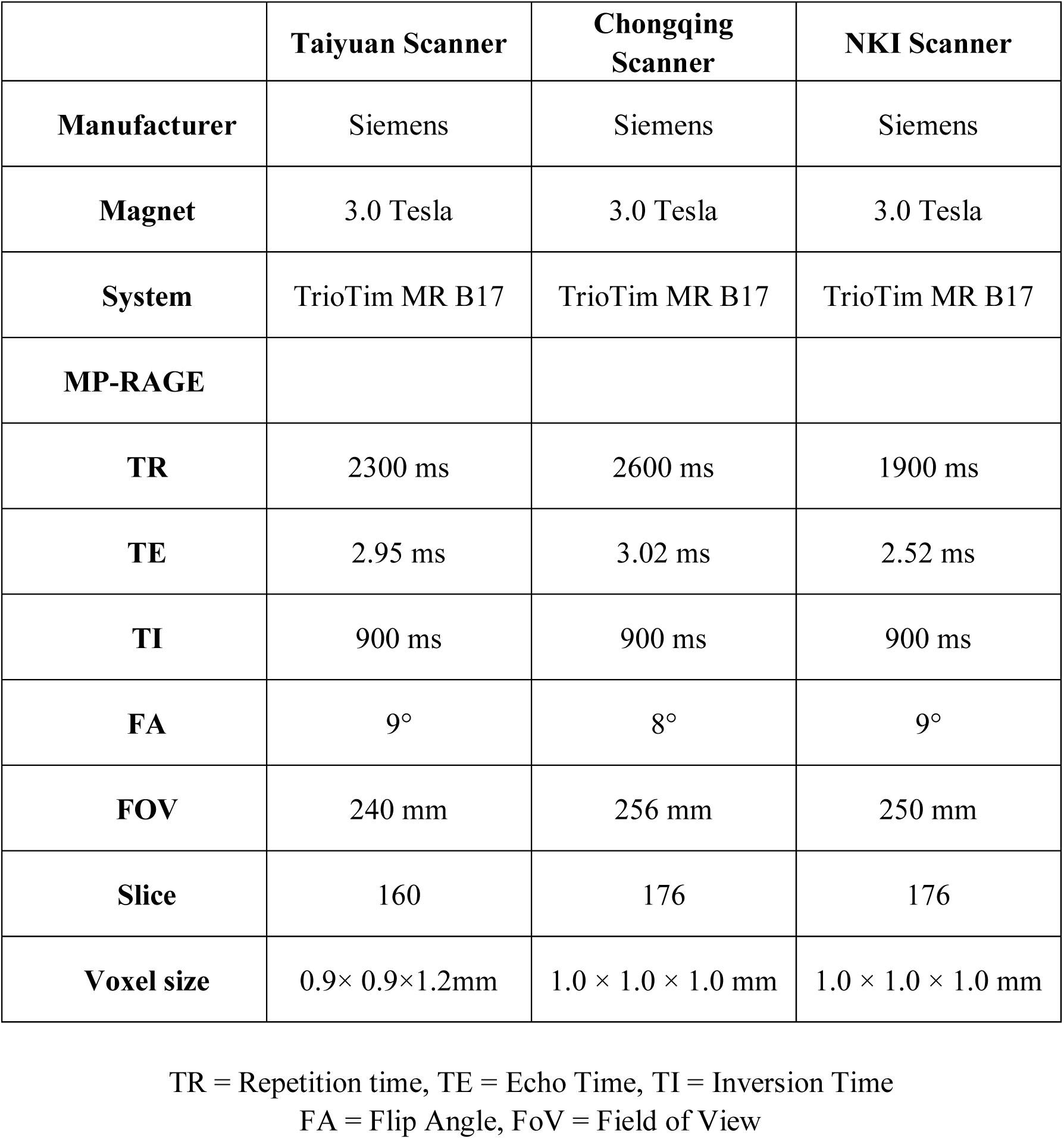
MRI scanning parameters at three imaging sites

|  | **Taiyuan Scanner** | **Chongqing Scanner** | **NKI Scanner** |
| --- | --- | --- | --- |
| **Manufacturer** | Siemens | Siemens | Siemens |
| **Magnet** | 3.0 Tesla | 3.0 Tesla | 3.0 Tesla |
| **System** | TrioTim MR B17 | TrioTim MR B17 | TrioTim MR B17 |
| **MP-RAGE** |  |  |  |
| **TR** | 2300 ms | 2600 ms | 1900 ms |
| **TE** | 2.95 ms | 3.02 ms | 2.52 ms |
| **TI** | 900 ms | 900 ms | 900 ms |
| **FA** | 9° | 8° | 9° |
| **FOV** | 240 mm | 256 mm | 250 mm |
| **Slice** | 160 | 176 | 176 |
| **Voxel size** | 0.9× 0.9×1.2mm | 1.0 × 1.0 × 1.0 mm | 1.0 × 1.0 × 1.0 mm |

TR = Repetition time, TE = Echo Time, TI = Inversion Time
FA = Flip Angle, FoV = Field of View

### Procedure

Brain template construction and growth chart modeling were completed using a standardized pipeline (Figure 7). To incorporate atlas information, we also performed a two-step protocol of image registration (steps 5-7). The procedure is as follows:

**Figure 7.**
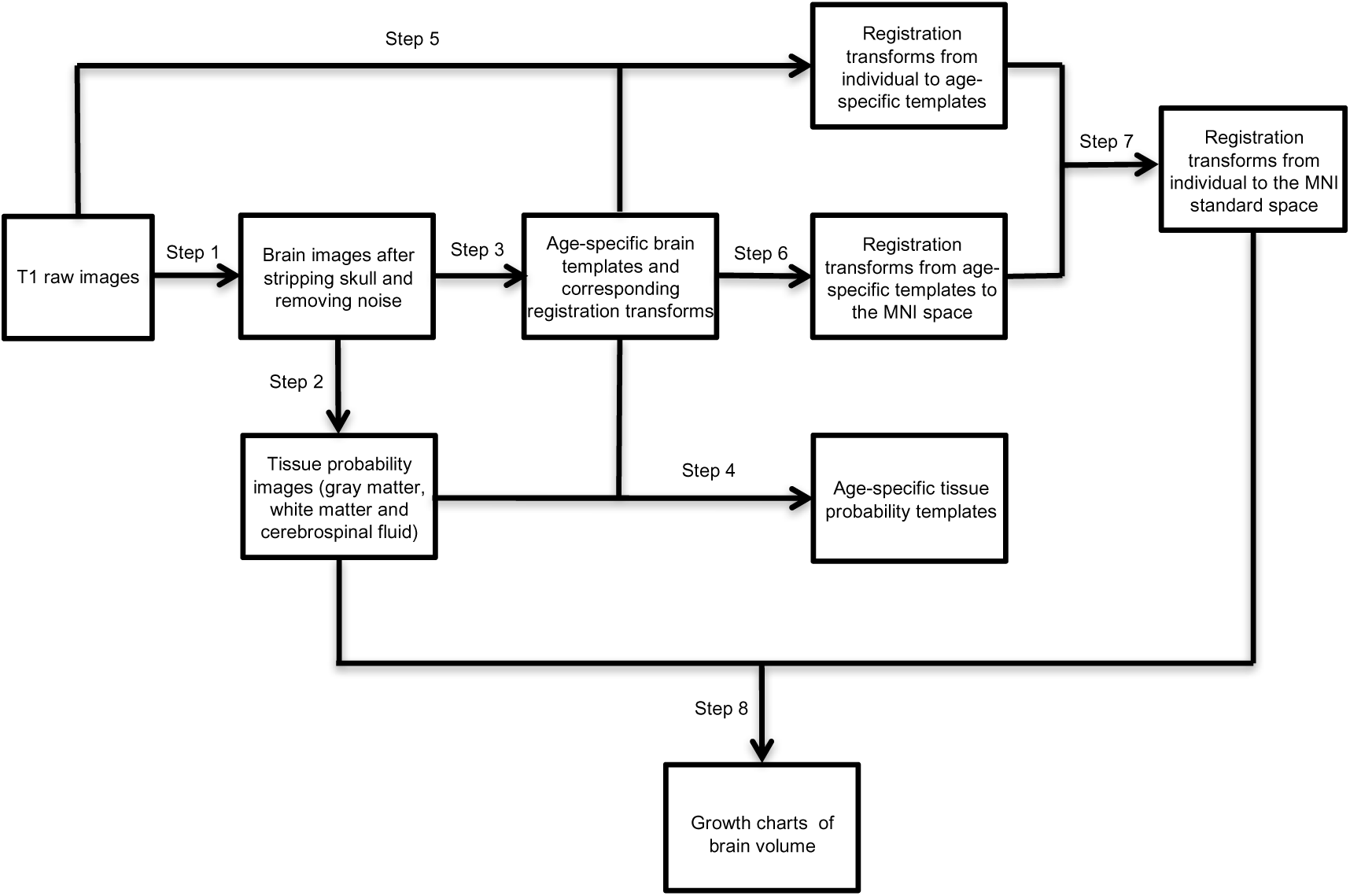
Standardized pipeline of brain template and growth chart estimation.

#### Quality check for T1 image and preprocessing: steps 1-2

**Step 1**. All individual high-resolution T1 images first underwent visual inspection to exclude images with substantial head motion and structural abnormalities. After initial quality control, the T1 images were entered into the *volBrain* pipeline (http://volbrain.upv.es/)55, which is an online program to remove image noise, intensity normalization and skull stripping. More specifically, noise artifacts, especially those showing spatially varying profiles, were suppressed using a highly effective filter with spatially adaptive nonlocal means (SANLM)^61, 62^. Initial image inhomogeneity correction was performed using N4 bias correction^63^. Next, the individual images were transformed to the MNI152 standard template space using advanced normalization tools (ANTs) with a linear transformation^64^ and further processed by fine correction of image inhomogeneity to improve image quality^65^. A piecewise linear mapping of image intensity^66^ was performed to normalize the intensities of gray matter (GM), white matter (WM), and cerebrospinal fluid (CSF) to mean intensities of 150, 250 and 50, respectively. This procedure was implemented to eliminate the effect of site on MRI signals and to improve brain extraction and skull stripping. Brain extraction was implemented using the nonlocal intracranial cavity extraction (NICE) algorithm^67^, which is an evolution of brain extraction based on the nonlocal segmentation technique (BEaST)^68^. NICE was built based on a multi-atlas label fusion strategy and a library of segmentation priors (pediatric dataset: age = 24.8 ± 2.4 months; normal adult dataset: age = 24 - 75 years) required for intracranial cavity labeling and has been demonstrated to outperform other skull stripping methods.

The above brain extraction was visually inspected to detect misclassification of tissues. If brain tissue was excluded from the segmentation, the brain mask was manually edited to ensure the quality of the brain extraction. To further check image quality, the signal-to-noise ratio (SNR), the contrast-to-noise ratio (CNR), the GM SNR and the WM SNR were computed for each image as described in reference^69^. Specifically, the SNR was calculated as the mean signal within the brain tissue divided by the standard deviation of the air signal. The GM SNR and the WM SNR were calculated as the SNR within the GM and WM tissue, respectively. The CNR was calculated as the mean GM intensity minus the mean WM intensity divided by the standard deviation of the air intensity. Any image with any of these quantitative metrics below the 1^st^ percentile was discarded. All the above steps were implemented as part of the Connectome Computation System (CCS) shared via GitHub^70^.

**Step 2**. To obtain probability tissue maps of individual brains, we segmented all individual brain images into GM, WM and CSF using the FSL FAST algorithm without settings of any prior probability maps^71^. Partial volume maps derived from FAST were used to estimate the proportion of a specific tissue within a voxel, measuring the tissue’s probability to construct tissue probability templates. Intracranial volume (ICV) was the number of all the voxels in the brain mask.

**Example FAST command for tissue classification:**

~~~
fast -n 3 -g -b -o <individual_pve> -p <input brain image>
~~~

#### Brain template construction: steps 3-4

**Step 3**. The brain template construction pipeline in ANTs was employed to build the pediatric brain templates. This pipeline requires no prior information for template construction^72^ and has been applied to the pediatric sample previously^73^. It is capable of extracting population-level representative images of the brain and other tissues such as the lungs^72–75^. Specifically, an optimal unbiased template was derived using the symmetric group-wise normalization (SyGN) algorithm in diffeomorphic space with respect to both shape and appearance^72^. SyGN first averages individual brains to obtain an initial brain template requiring no specific a priori information. A nonlinear iterative procedure of deformation was then performed as follows: 1) the optimal deformation was computed according to the initial mean template for each individual image; 2) a template to maximize the similarity metric between the template and individual images was performed using a gradient descent algorithm (only the template appearance is updated while the template shape and mappings are fixed); 3) the template shape was optimized; 4) the parameters derived from the above steps were subsequently updated, and a new template was generated as the reference mean image in step 1 for the next iteration. For a more stable template, we set the number of iterations to 10 for all age groups, taking into account that the algorithm usually converged after 3 to 5 iterations^72^.

We reconstructed the brain and skull templates separately and combined them into head templates for each age group. We chose cross-correlation as the similarity metric and Greedy B-spline SyGN as the transformation model for the brain registration, with shrinkage factors, smoothing factors and max iterations of 8*×*4*×*2*×*1, 3*×*2*×*1*×*0 and 100*×*70*×*50*×*10, respectively. To achieve comparable space without any shape changes, each brain template was rigidly transformed into the MNI152 template space using ANTs.

**Example ANTs command for template construction:**

~~~
antsMultivariateTemplateConstruction2.sh -d 3 -o
<output_brain_template> -i 10 -g 0.25 -c 4 -k 1 -w 1 -f 8×4×2×1 -s 3×2×1×0 -q 100×70×50×10 -n 1 -r 1 -l 1 -m CC[2] –t BSplineSyN[0.1,26,0]
<input_brain_images>
(output_brain_template is the name of output file in this step, defined by users.)
~~~

**Step 4**. The affine matrices (linear) and deformation transforms (nonlinear) generated in step 3 were combined and applied to the individual tissue segmentation images. Then tissue probability templates were constructed for each age group; head templates were also reconstructed by combining brain templates and skull templates for each age group.

**Example ANTs commands for applying registration transforms:**

**Apply registration transform files generated in step 3 on individual brain tissue probability files:**

~~~
antsApplyTransforms -d 3 -i <individual_pve> -o <tmp_pve> -r <template_generated_in_step3> -t <Warpfile> -t <Affinefile>
(Warpfile refers to the nonlinear deformation file for each subject, Affinefile refers to the linear affine transformation file for each subject, both files can be found in the template output directory set in step 3, Warpfile is denoted as (subject)*1Warp.nii.gz while Affinefile denoted as (subject)*GenericAffine.mat)
AverageImages 3 <output_AVG> 0 sub01_pve sub02_pve … subN_pve
(output_AVG is the name of output file in this step, defined by users. SubN_pve refers to the output tissue probability files generated from step 2.)
antsApplyTransforms -d 3 --float 1 --verbose 1 –i <output_AVG> -o <output_tissue_template> -t [<Afffile>,1] -t <Warpfile> -t <Warpfile> -t <Warpfile> -t <Warpfile> -r <output_brain_template>
(output_tissue_template is the name of output file in this step, defined by users. Warpfile refers to the averaged deformation file for template, Affinefile refers to the averaged affine transformation file for template, both files can be found in the template output directory set in step 3, Warpfile is denoted as *template0Warp.nii.gz while Affinefile denoted as *template0GenericAffine.mat)
~~~

#### Two-step registration from individual images to MNI152 space: steps 5-7

**Step 5**. All individual images were registered to the appropriate template using ANTs. To explore the extent to which registration errors affect tissue volume estimation, registration to mismatched templates were also performed for growth chart modeling, that is, images from the CCNP sample were registered to the age appropriate NKI brain template.

**Example ANTs command for registration:**

~~~
antsRegistrationSyN.sh -d 3 -f <fixed_Individua_Image> -m <AST> -o <reg2AST>
~~~

(AST refers to Age Specific Template.)

**Step 6**. Age-specific templates were registered to the MNI152 template using ANTs, the most commonly used standard space coordinate system for overlaying brain atlas and parcellation information.

**Example ANTs command for registration:**

~~~
antsRegistrationSyN.sh -d 3 -f <AST> -m MNI152.nii.gz -o <reg2MNI>
~~~

**(AST refers to Age Specific Template.)**

**Step 7**. Combining the registered transforms generated in steps 5 and 6, the individual brain images were warped to the MNI152 template for the convenience of delivering brain atlas information. This two-step registration is designed to avoid systematic bias induced by direct registration of individual brain images to mismatched age- and Chinese-specific brain templates while maintaining the integrity of the brain atlas information as much as possible.

To better demonstrate the application of age-specific templates and elucidate brain trajectories at more detailed structural levels, brain lobes and area-level parcels were delivered from standard MNI template into individual space at this step, followed by parcel volume extraction which were passed into growth chart modeling subsequently.

**Example ANTs command for combining transforms generated from steps 5 and 6 (take brain lobe mask registration for instance):**

~~~
antsApplyTransforms -d 3 -n NearestNeighbor -i <lobe_mask> -o <ASToutput> -r <AST> -t [<reg2MNI_affine>,1] -t <reg2MNI_Inwarp>
(This command registers the mask file defined on MNI152 template to Age Specific Template. reg2MNI_affine refers to the affine files generated in step 6, reg2MNI_Inwarp refers to the inverse warp files generated in step 6, if the fixing image in step 6 was set to MNI152 template and moving image set to AST, then the warp file should be applied here instead of inverse warp files.)
antsApplyTransforms -d 3 -n NearestNeighbor -i <ASToutput> -o <Individual_lobe_mask> -r <Individual_image> -t [<reg2AST_affine>,1] -t <reg2AST_Inwarp>
(This command registers the mask file generated from the above command to individual image. reg2AST_affine refers to the affine files generated in step 5, reg2AST_Inwarp refers to the inverse warp files generated in step 5, if the fixing image in step 5 was set to Age Specific template and moving image set to individual image, then the warp file should be applied here instead of inverse warp files.)
~~~

#### Growth chart modeling

**Step 8**. The dynamic developmental process was modeled with growth charts of different brain tissues to highlight the need for age-specific brain templates. We extracted ICV and its three tissue components (GM, WM, CSF) using FAST in FSL^71^. To obtain corresponding quantification at the lobar level (frontal, temporal, parietal and occipital) and regional levels, we registered the lobe and area parcels from the MNI152 template to the current age-specific templates and then to the individual space; lobe and regional level GM volumes were extracted by multiplying GM probabilities and total volumes within individual lobe parcels.

Quantile regression was employed to build brain growth charts^76^. We chose the LMS method of centile estimation to construct the growth curves of brain sizes and volumes. Specifically, this method summarizes the age-related nonlinear distribution of the measurement of interest by 3 curves, representing the median (M), coefficient of variation (S), and skewness (L) of the distribution. These curves can be fitted as cubic splines by nonlinear regression, where the smoothing extent required can be expressed in terms of smoothing parameters or equivalent degrees of freedom. The above analysis was performed using GAMLSS implemented in R (version 3.4.3)^56^. Two models have been conducted to explore developmental trajectories. In one model, volume data of all subjects was utilized for growth charts modeling, while in the other, growth curves were modeled separately for boys and girls. This analytic strategy has been employed by the World Health Organization (WHO) and Centers for Disease Control and Prevention (CDC) to delineate growth charts of height and weight for children^77–79^.

**Commands for modeling Growth Charts (R):**

~~~
library(gamlss) library(gamlss.dist)
GCdata <- read.table(“DATAset”,header = TRUE)
GCmodel <- lms(TissueVolume, age, data=GCdata, method.pb=“GAIC”, k=5)
Age_predict <- seq(6,18,0.25)
centiles(GCmodel, GCdata$age, cent=c(5,25,50,75,95), legend=FALSE, ylab=“GCmodel”, xlab=“Age”, pch=“o”, lwd.centiles=c(2.5,2.5,4,2.5,2.5))
~~~

(DATAset refers to the tissue developmental data generated in the previous step, which comprises one variable named ‘TissueVolume’ referring to the volume of brain tissue or parcels while another variable named ‘age’ referring to subject age.)

### Timing

Step 1 takes 10-20 minutes per subject. Step 2 takes approximately 5 minutes using a computer with a Xeon E5 2GHz CPU. Template construction in step 3 takes considerable time, depending on sample size and number of iterations. For instance, the 11-year-old template built from 108 images with 10 iterations took 17 hours 7 minutes using ANTs. Step 4 should take about 5-10 minutes depending on sample size. In steps 5-7, the most time-consuming operation is ANTs registration (about 50 minutes to register to the age-specific template per subject). (Total time was about 20 hours for the 11-year-old group.) All data processing was performed on a cluster server with 24 nodes and 300 CPU cores at IPCAS, which processed the registration computation in parallel.

### Validation of Template Use

Two new pediatric neuroimaging datasets from Weifang Medical University^80^ and Zhejiang University^81, 82^, including 84 structural MRI scans, were employed to validate the necessity of constructing age- and ethnicity-matched MRI templates using brain deformation cost function (see age, sex and scanning protocols in Tables 5 and 6).

**Table 5.** Gender composition in validation samples with one-year increment

| age range | Weifang Site |  | Zhejiang Site |  | male | total |
| --- | --- | --- | --- | --- | --- | --- |
|  | male | total | male | total |  |  |
| **7.0-7.9** | 2 | 8 |  |  | 2 | 8 |
| **8.0-8.9** | 3 | 3 | 0 | 1 | 3 | 4 |
| **9.0-9.9** | 2 | 7 | 5 | 14 | 7 | 21 |
| **10.0-10.9** | 6 | 12 | 17 | 27 | 23 | 39 |
| **11.0-11.9** | 3 | 4 | 6 | 8 | 9 | 12 |
| **total** | 16 | 34 | 28 | 50 | 44 | 84 |

**Table 6.** MRI scanning parameters of validation samples

|  | **Weifang Site** | **Zhejiang Site** |
| --- | --- | --- |
| **Manufacturer** | Siemens | Philips |
| **Magnet** | 3.0 Tesla | 3.0 Tesla |
| **System** | TrioTim MR B17 | Achieva |
| **TR** | 2530 ms | 30 ms |
| **TE** | 3.37 ms | 5 ms |
| **TI** | 1100 ms |  |
| **FA** | 7° | 15° |
| **FOV** | 256 mm | 230 mm |
| **Slice** | 128 | 150 |
| **Voxel size** | 1.0 × 1.0 × 1.33 mm | 0.41 × 0.41 × 1.0mm |

TR = Repetition time, TE = Echo Time, TI = Inversion Time FA = Flip Angle, FoV = Field of View

### Procedure

The brain templates were validated by a standardized pipeline, the details are as follows:

**Validation-Step 1.** The validation data were subjected to the same preprocessing pipeline as the template data described in last section, step 1.

**Validation-Step 2.** Each individual T1-weighted (T1w) brain image, following denoising and skull stripping, was fed into ANTs for further registration to each brain template. For each individual, 11 registrations were performed with our Chinese age-specific pediatric (IPCAS) MRI templates, and 11 registrations with the NKI pediatric MRI templates.

**Validation-Step 3.** We calculated the warping distance at each voxel for deformation registration and then averaged the values across all brain voxels to represent the extent of individual deformation. The individual warp values were transferred into Z scores for inter-subject group analysis. For each age-specific template, a template age was also obtained by averaging all subjects’ ages within the group, which later subtract the age of the subject of each scan in the validation group to represent the age difference between the target template and the source individual brain image. An age difference of zero indicates a perfect match between the age of the template and the age of the individual, while a negative age difference indicates that the individual is older than the template age, and a positive value indicates that the individual is younger than the template age. As the age span in the validation group ranged from 7 to 12 years, the resulting age differences ranged from −5 to 10 years. Generalized Additive Models for Location Scale and Shape (GAMLSS) were finally applied to model changes in the registration warp curve with age differences. We expected to observe an age effect for registration deformation, that is, more age mismatch between the target template and the source individual would result in more registration deformations and registration costs, i.e., less efficient registration.

### Comparison of Brain Growth Charts (CCNP vs. eNKI)

To quantitatively estimate the diversity of growth charts attributed to ethnicity, the normalized variance (NV) was calculated across 400 brain parcel units in MNI space^47^ contrasting CCNP and eNKI samples, with the NV values calculated as follows:

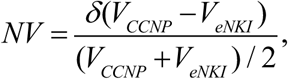

where V is a vector referring to the parcel unit volume at every age point estimated in the last step; the standard deviation of the differences between the two samples was calculated to characterize the degree of chart shape dispersion across different ages. To exclude potential confounding, it was normalized by the mean parcel volume. Large NV values indicate diversity while small values indicate the growth curves share similar shapes (Figure 8).

**Figure 8.**
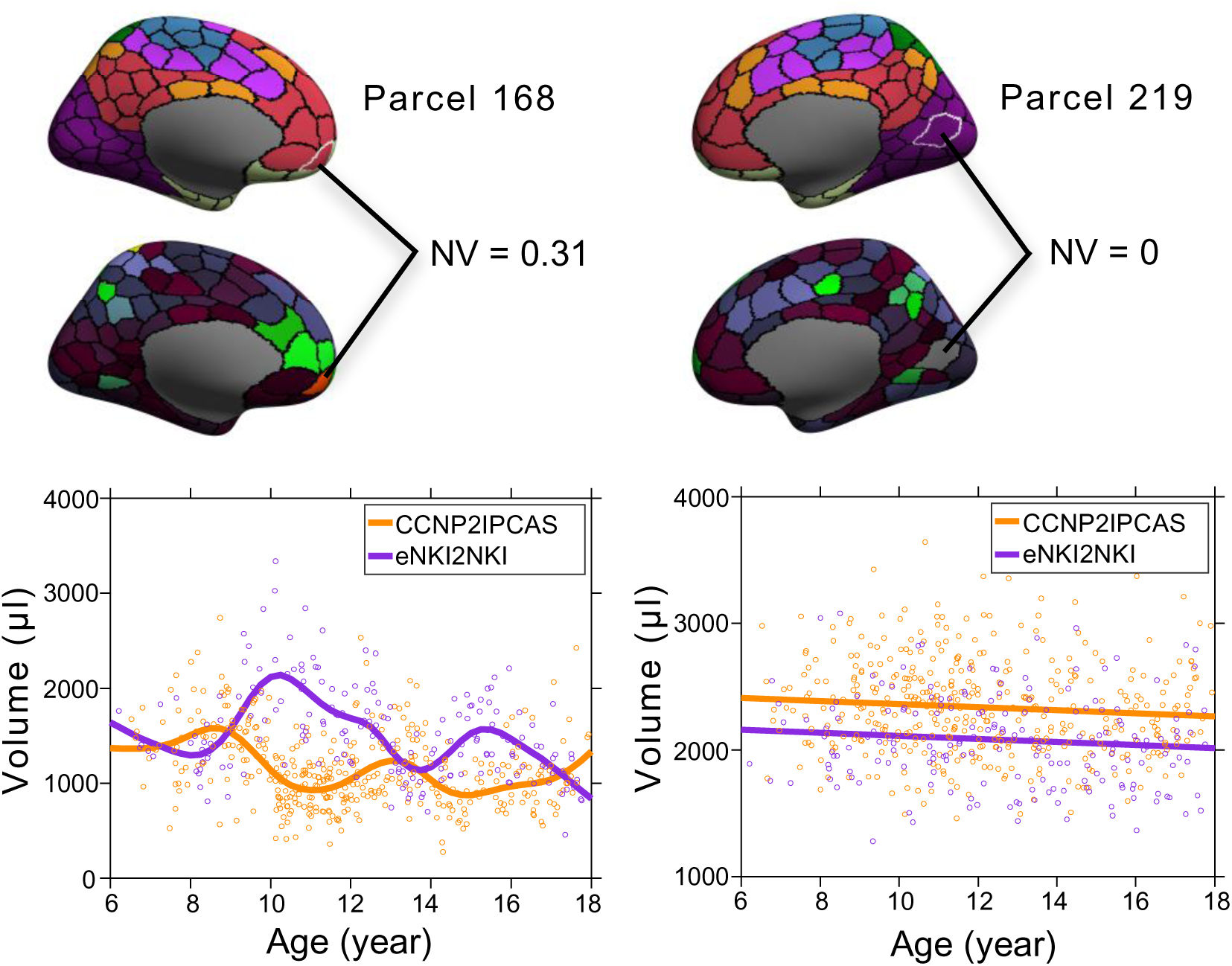
Two parcel units exhibited the most similar and different developmental patterns. Upper panel shows the locations of the two parcels with largest and smallest NV value, and lower panel shows the corresponding curves of the two parcels.

### Anticipated Results

This protocol produced standard brain and tissue probability templates, and growth charts for brain tissue and areas. The axial slices of age-specific templates are illustrated in Figure 1. The upper four rows were built from the CCNP sample while the lower four rows were constructed from the eNKI sample. Ages are displayed across columns, from ages 6-7 to 17. Clear differences in tissue spatial profiles across childhood and adolescence are observable, and ethnic differences in brain shape can be appreciated.

Growth charts including global metrics like intracranial volume (ICV), GM volume, WM volume and CSF are also displayed in Figure 3 for CCNP and eNKI samples with all subjects combined and separately for males and females in sFigure 1.

At a more refined scale, growth charts of regional brain volumes were compared. The similarity of trajectories between CCNP and eNKI samples was estimated for each area and depicted in Figure 5. Large differences were mostly observed in the association cortex while the primary cortex exhibited similar developmental trajectories. To better define the distribution of NVs among hierarchical brain networks, the bar graphs of regional NVs is also shown in Figure 5, with colors indicating the 7 large scale brain networks and left and right hemispheres shown separately. Overall, the left hemisphere demonstrated greater diversity, specifically in the default and dorsal attention networks, although the right hemisphere frontal-parietal network varied the most between ethnicities. The regional trajectories of maximum and minimum NVs in the left hemisphere are shown in Figure 8, with an absolute opposite pattern clearly revealed for the area with the largest NV value. The trajectories of areas with small NV values were almost identical across childhood and adolescence. Compared with the variety in mesoscopic brain areas, developmental trajectories at the level of brain lobes exhibited more similar patterns (sFigure 2).

## Funding

This work was supported in part by the Natural Science Foundation of China (81220108014), the National Basic Research (973) Program (2015CB351702), the China - Netherlands CAS-NWO Programme (153111KYSB20160020), Beijing Municipal Science and Tech Commission (Z161100002616023, Z171100000117012), the Major Project of National Social Science Foundation of China (14ZDB161), the National R&D Infrastructure and Facility Development Program of China, Fundamental Science Data Sharing Platform (DKA2017-12-02-21), and Guangxi BaGui Scholarship (201621). BTTY is funded by the Singapore National Research Foundation (NRF) Fellowship (Class of 2017).

## Conflict of Interest Statement

The authors declare that the research was conducted in the absence of any commercial or financial relationships that could be construed as a potential conflict of interest.

## Acknowledgments

The authors thank Dr. Arno Klein from Child Mind Institute and Dr. Zhi Yang from Shanghai Mental Health Center for their highly valuable comments on templates validation and brain morphological metric quantifications. We would like to thank all parents and children participating in this study as well as all the support from schools and community.

