## Supplemental Materials for "Generating Templates and Growth Charts for School-Aged Brain Development"

**SFigure 1 Developmental curves of Intracranial volume (ICV), gray matter (GM), white matter (WM) and cerebral spinal fluid (CSF).**

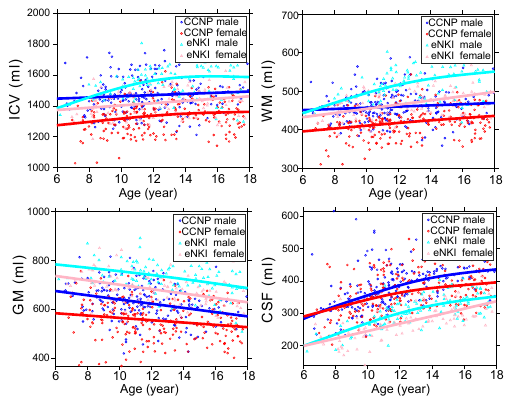

**SFigure 2 Developmental curves of brain lobe volume.**

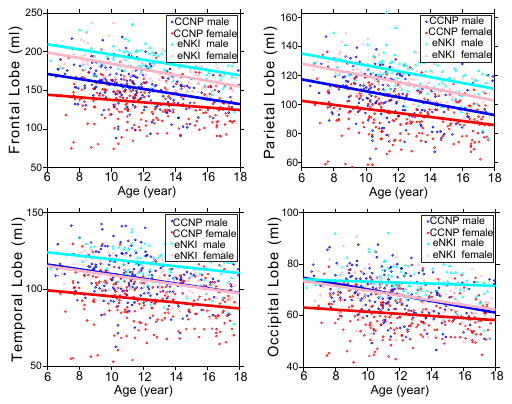
